# Identification of structurally diverse FSP1 inhibitors that sensitize cancer cells to ferroptosis

**DOI:** 10.1101/2022.12.14.520445

**Authors:** Joseph M. Hendricks, Cody Doubravsky, Eddie Wehri, Zhipeng Li, Melissa A. Roberts, Kirandeep Deol, Mike Lange, Irene Lasheras-Otero, S.J. Dixon, Kirill Bersuker, Julia Schaletzky, James A. Olzmann

## Abstract

Ferroptosis is a regulated form of cell death associated with the iron-dependent accumulation of lipid peroxides. Inducing ferroptosis is a promising approach to treat therapy resistant cancer. Ferroptosis suppressor protein 1 (FSP1) promotes ferroptosis resistance in cancer by generating the antioxidant form of coenzyme Q10 (CoQ). Despite the important role of FSP1, few molecular tools exist that target the CoQ-FSP1 pathway. Exploiting a series of chemical screens, we identify several structurally diverse FSP1 inhibitors. The most potent of these compounds, ferroptosis sensitizer 1 (FSEN1), is an uncompetitive inhibitor that acts selectively through on target inhibition of FSP1 to sensitize cancer cells to ferroptosis. Furthermore, a synthetic lethality screen reveals that FSEN1 synergizes with endoperoxide-containing ferroptosis inducers, including dihydroartemisinin, to trigger ferroptosis. These results provide new tools that catalyze the exploration of FSP1 as a therapeutic target and highlight the value of combinatorial therapeutic regimes targeting FSP1 and additional ferroptosis inducers.

## INTRODUCTION

Ferroptosis is an iron-dependent, non-apoptotic form of regulated cell death that is associated with the lethal accumulation of oxidatively damaged phospholipids (e.g., lipid peroxides)^1,2^. As the ultimate effectors, phospholipid hydroperoxides, or their breakdown products, mediate membrane rupture and ferroptotic cell death^1,2^. Cellular ferroptosis sensitivity is determined by a variety of metabolic processes, including pathways that regulate labile iron pools and the generation of reactive oxygen species (ROS), the addition and removal of oxidation-sensitive polyunsaturated fatty acids (PUFAs) to and from phospholipids, and ferroptosis defense systems that suppress the accumulation of lipid peroxides^1,2^. These ferroptosis defense systems function through two mechanisms, the conversion of lipid peroxides into non-toxic lipid alcohols by the glutathione (GSH) peroxidase GPX4^3^ and the generation of endogenous antioxidants that prevent lipid radical propagation, such as the generation of the reduced form of coenzyme Q10 (CoQ) (i.e., ubiquinol) by ferroptosis suppressor protein 1 (FSP1)^4,5^ and dihydroorotate dehydrogenase (DHODH)^6^, tetrabiohydropterin by GTP cyclohydroxylase-1 (GCH1)^7,8^, and hydropersulfides by enzymatic and non-enzymatic pathways^9,10^.

Ferroptosis serves as a natural mechanism to restrict cancer proliferation and survival that is engaged by tumor suppressors. For example, the expression of the system x_c_^-^ antiporter subunit SLC7A11, required for cystine uptake and GSH synthesis, is inhibited by the tumor suppressors BAP1^11^, p53^12^, and Kelch-like ECH-associated protein 1 (KEAP1)^13^. Inactivating mutations in these tumor suppressors upregulate SLC7A11 and other genes that promote ferroptosis resistance^14^. While oncogene-driven reprogramming of cancer cell metabolism addresses the amplified cellular demands for nutrients and energy, the accompanying increases in ROS, and a consequent overreliance upon ferroptosis defensive systems for survival, yield potential therapeutic vulnerabilities^14–16^. Indeed, inhibition of the GSH-GPX4 pathway is effective in killing many cancer cells *in vitro* and in reducing tumor growth in preclinical models of therapy resistant cancers such as pancreatic ductal adenocarcinoma (PDAC)^17,18^, clear cell and chromophobe renal carcinomas^19,20^, triple-negative breast cancer^21^, MYCN-amplified neuroblastoma^22–25^, and drug-resistant persister cancer cells that give rise to relapse^26,27^. Despite these promising findings, some cancer cells are resistant to inhibition of the GSH-GPX4 pathway due to compensation by parallel ferroptosis defense systems. However, in contrast to the GSH-GPX4 pathway^28^, few molecular tools are available that target the other ferroptosis defense systems such as the CoQ-FSP1 pathway.

FSP1 is an important ferroptosis resistance factor in cancer that compensates for the loss of GPX4 by mediating NAD(P)H-dependent reduction of ubiquinone (oxidized form of CoQ) to ubiquinol (reduced form of CoQ), which in turn acts as a lipophilic antioxidant to prevent lipid peroxidation propagation^4,5^. FSP1 expression is amplified in ferroptosis resistant non-small cell lung cancer and its expression correlates with poor patient prognosis^29^. Genetic disruption of FSP1 sensitizes cancer cells to ferroptosis and impairs tumor growth in a model of KEAP1-deficient lung cancer^4,29^. The tumor suppressor KEAP1 is an adaptor for the cullin-3 E3 ubiquitin-protein ligase that mediates the proteasomal degradation of nuclear factor erythroid 2-related factor 2 (NRF2), a master transcription factor that regulates the expression of an antioxidant gene program^30,31^. Thus, inhibitors of FSP1 have potential therapeutic value as anti-cancer monotherapies or when used in combination with ferroptosis inducers such as compounds targeting the GSH-GPX4 pathway. Consistent with this possibility, an inhibitor of FSP1 (iFSP1) sensitized multiple cancer cell lines to ferroptosis triggered by GPX4 inhibition and radiotherapy^5,29^. iFSP1 provides an important proof of concept, but additional structurally distinct FSP1 inhibitors with improved pharmacokinetic and pharmacodynamic properties are critically needed to fully explore the potential of FSP1 as a therapeutic target.

To address the unmet need for FSP1 inhibitors, we conducted a series of small molecule screens that leverage an *in vitro* assay of FSP1 CoQ oxidoreductase activity and an orthogonal cell-based assay of FSP1-dependent ferroptosis suppression. These screens identified multiple structurally distinct small molecules that directly inhibit FSP1 CoQ oxidoreductase activity *in vitro* and sensitize cancer cells to ferroptosis. Moreover, a second screen of FDA-approved and bioactive compounds identified a synergistic relationship of FSP1 inhibition with the endoperoxide-containing drug dihydroartemisinin (DHA) in inducing ferroptosis. Our study provides new molecular tools for the characterization of FSP1 as an anti-cancer therapeutic target and demonstrates the utility of combinatorial treatment regimens targeting FSP1 together with other ferroptosis defense pathways.

## RESULTS

### Chemical library screen identifies small molecule inhibitors of FSP1 activity

To identify small molecule inhibitors of FSP1, we conducted a chemical screen employing an *in vitro* assay of FSP1 activity (**Figure 1A,B**). This assay exploits the change in 355 nm absorbance as NADH is oxidized to NAD+ during the reduction of CoQ1 by recombinant FSP1 (**Figure 1B,C**). As anticipated, the addition of FSP1 to the reaction mix resulted in a decrease in absorbance over time (**Figure 1C**). Moreover, the decrease in absorbance was blocked in a dose-dependent manner by iFSP1 (IC50 of 4 µM) (**Figure 1C, Figure S1A**). These data demonstrate that iFSP1 is a direct FSP1 inhibitor and validates our activity assay as a method to identify FSP1 inhibitors.

**Figure 1.**
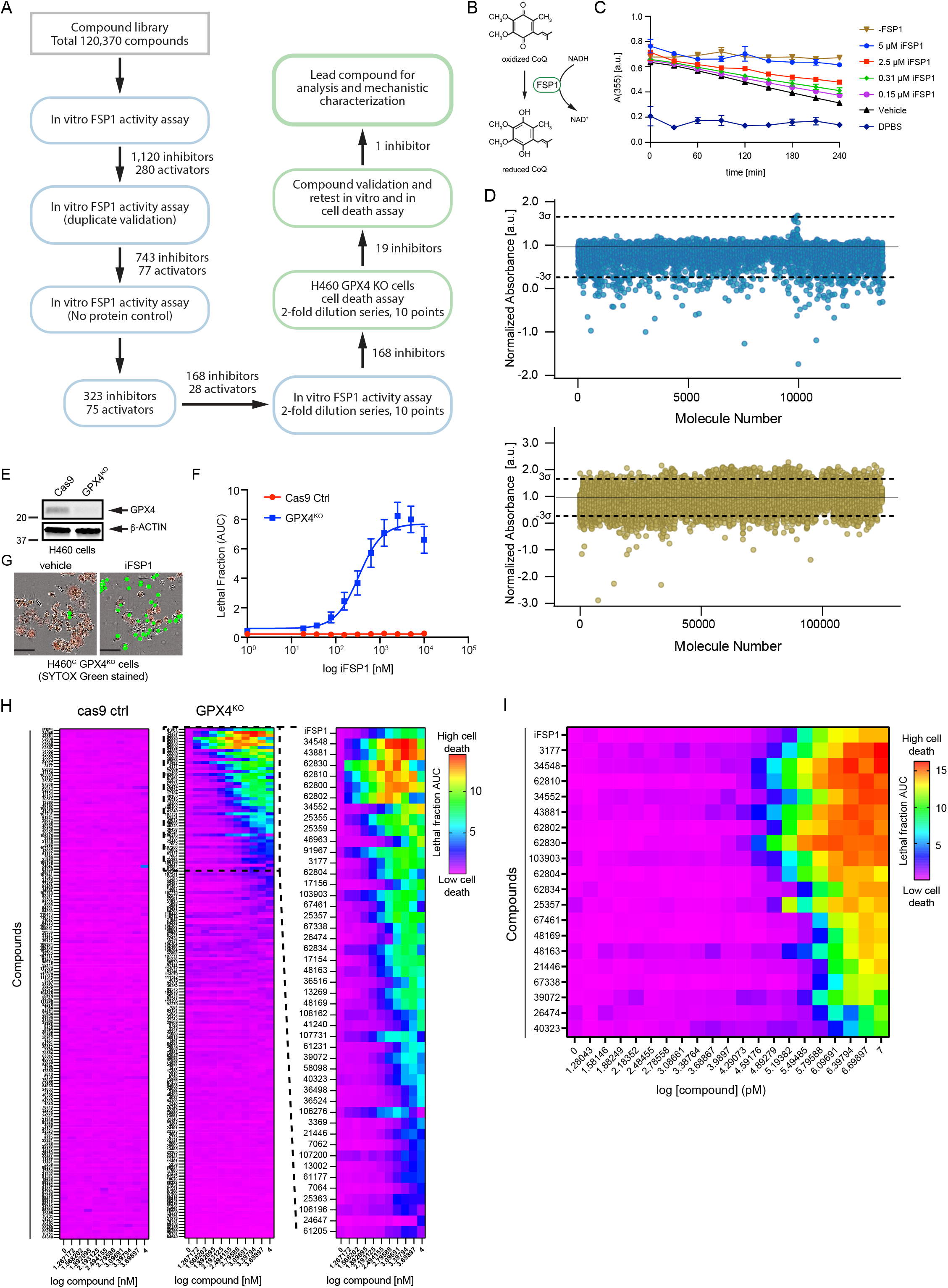
Small molecule screens identify FSP1 inhibitors. **A)** Flow chart of the chemical screens and experimental validation employed to identify FSP1 inhibitors. **B)** Schematic of the *in vitro* activity assay using purified recombinant FSP1. **C)** FSP1 activity was measured using NADH absorbance in the presence of increase amounts of iFSP1. Mean ± SEM of three technical replicates. **D)** Scatter plots of 15,000 antibacterial compounds and 100,000 diverse + 5,370 FDA and Bioactive compounds assayed with *in vitro* absorbance-based assay of FSP1 activity. “Hits” are defined by a normalized absorbance value of <0.267. Dashed lines represent SD of 3-sigma cutoff defined by activity range of vehicle and no protein control. **E)** Western blot analysis of H460^C^ Cas9 and GPX4^KO^ cells. **F)** Representative images of H460 GPX4^KO^ cells treated with vehicle and iFSP1. Dead cells are marked with SYTOX Green. Scale bar represents 50 µm. **G)** Dose response of iFSP1 induced cell-death in H460^C^ Cas9 and GPX4^KO^ cells. Lethal fraction was calculated by IncuCyte quantification of the ratio of dead cells over the total amount of cells over 24 hr. The mean ± SEM bars represent three biological replicates. **H)** Lethal fraction (AUC) was calculated in H460^C^ Cas9 and GPX4^KO^ cells incubated with increasing doses of 168 small molecules that inhibited FSP1 *in vitro*. The heat map reflects three biological replicates. **I)** Lethal fraction was calculated H460^C^ GPX4^KO^ cells incubated with increasing doses of the most potent 19 FSP1 inhibitors and iFSP1. Resupplied, validated small molecules were used. The heat map reflects three biological replicates.

In the primary screen, the effect of 120,370 small molecules on FSP1 activity was analyzed using the *in vitro* FSP1 activity assay (**Figure 1A,D**). This screen identified 1,120 candidate FSP1 inhibitors and 660 candidate activators based on a 0.264 normalized absorbance threshold value (**Figure 1D**). Duplicate analyses of the candidate inhibitors and activators were performed for validation (**Figure S1B)**. A control lacking FSP1 protein was included to identify small molecules that altered absorbance independently of FSP1 (**Figure S1C**). This series of validation steps yielded 323 inhibitors and 75 activators. Finally, triplicate 10-point dose response analyses were performed to determine the *in vitro* potency of 168 selected of the FSP1 inhibitors. 26 of these compounds have a lower IC_50_ than iFSP1 (< 4 µM) and 11 compounds have an IC_50_ below 100 nM.

### Small molecule inhibitors of FSP1 trigger cell death in a cancer cell model

To determine whether the 168 candidate FSP1 inhibitors are able to inhibit FSP1 in cells, we developed an orthogonal cell-based assay of FSP1 activity. This assay uses NCI-H460 KEAP1 mutant lung cancer cells expressing mCherry (H460^C^) in which GPX4 was knocked out using CRISPR-Cas9 (H460^C^ GPX4^KO^) or which expressed Cas9 as a control (H460^C^ Cas9) (**Figure 1E**). mCherry was used as a live cell marker, which together with the SYTOX Green cell death marker can be used to calculate the fraction of dead cells (i.e., lethal fraction^32,33^). To validate this assay, H460^C^ Cas9 cells and H460^C^ GPX4^KO^ cells were treated with iFSP1 (**Figure 1F,G**). iFSP1 selectively triggered cell death in H460^C^ GPX4^KO^ cells, but not in the H460^C^ Cas9 cells (**Figure 1F,G**). These data demonstrate that this assay can be used to characterize the ability of FSP1 inhibitors to inhibit FSP1 and induce ferroptosis in cancer cells.

Employing this assay, we analyzed the amount of cell death induced by 168 FSP1 inhibitors (**Figure 1A,D**) in the H460^C^ Cas9 and H460^C^ GPX4^KO^ cells **(Figure 1H**). H460^C^ Cas9 cells were included to identify any small molecules that kill cells through a ferroptosis-independent mechanism. Triplicate 10-point dose response analyses were performed, and cell death was measured using fluorescence time-lapse imaging **(Figure 1H**). ∼50 of the FSP1 inhibitors induced cell death in the H460^C^ GPX4^KO^ cells (**Figure 1H**), but not the H460^C^ Cas9 cells, indicating that these compounds are synthetic lethal with GPX4^KO^ and are not generally toxic to cells.

19 of the most potent compounds were tested again using validated compounds (>95% purity) in 20-point dose response analyses in both the *in vitro* (**Figure S2 and S3**) and cell-based FSP1 assays (**Figure 1I**). As observed in our primary and follow up screens, these 19 compounds directly inhibited purified FSP1 *in vitro* (**Figure S2 and S3**) and triggered cell death in the H460^C^ GPX4^KO^ cells, but not the H460^C^ Cas9 cells (**Figure 1I**). We named these validated FSP1 inhibitors – ferroptosis sensitizer 1-19 (FSEN1-19). The structures of FSEN1-19, their IC50 for inhibition of purified FSP1 activity, and their EC50 for triggering cell death in H460^C^ GPX4^KO^ cells are shown in **Figure 2**. These compounds can be organized into seven groups of structurally related compounds. The largest group of compounds (Group 1, red box) share a disubstituted [1,2,4]triazolo-thiazole core scaffold structure (**Figure 2**).

**Figure 2.**
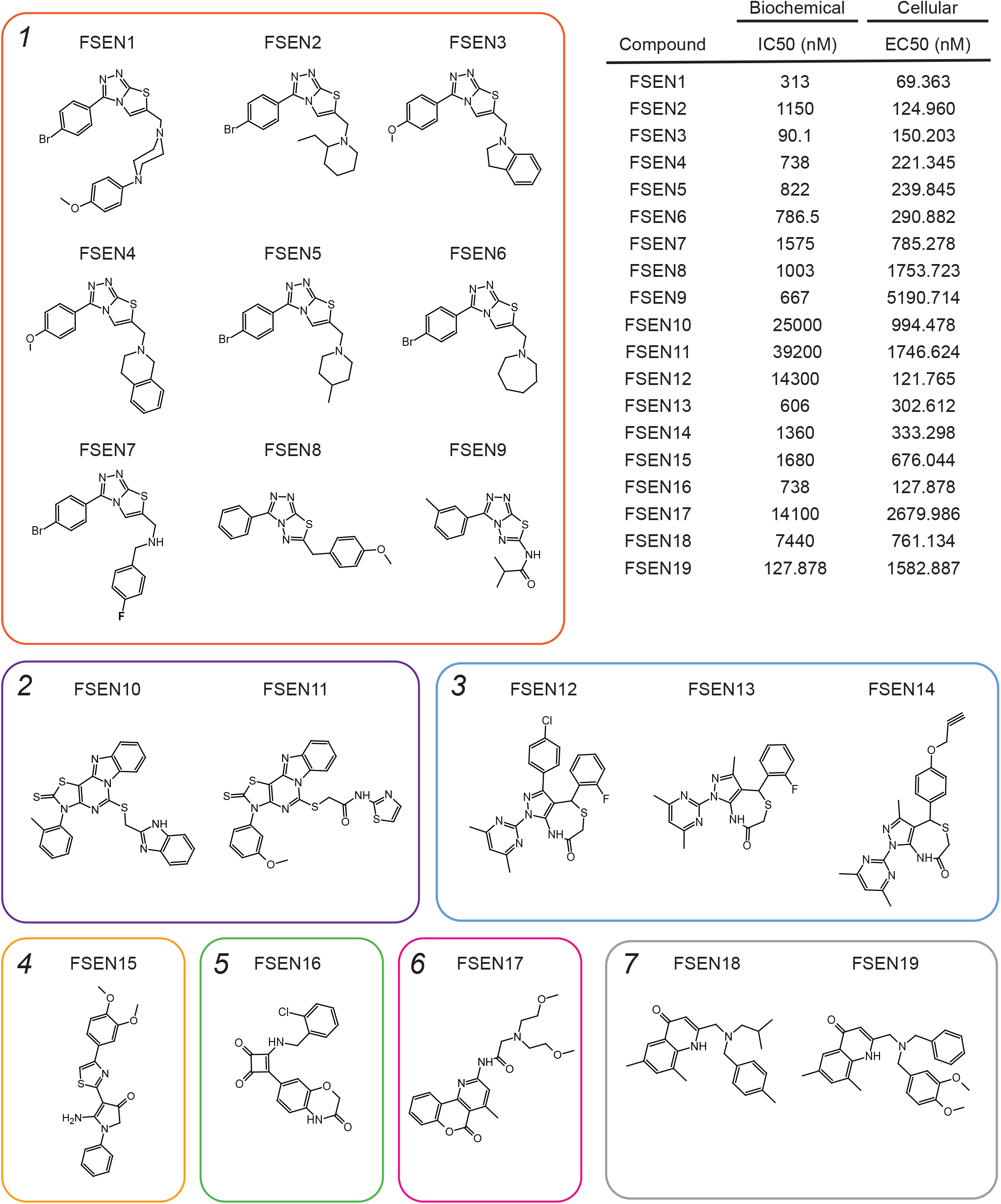
Multiple structurally distinct FSP1 inhibitor scaffolds. The structures of FSEN1-19 are shown with their calculated IC50 values from end-point assays measuring inhibition of FSP1 activity *in vitro* (n=2) and their calculated EC50 values from lethal fraction (AUC) quantification in H460^C^ GPX4^KO^ cells (n=3).

### FSEN1 is an uncompetitive inhibitor of FSP1

Amongst the newly identified FSP1 inhibitors, FSEN1 exhibited the highest potency in triggering cell death in the H460^C^ GPX4^KO^ cells (EC50 = 69.363 nM) (**Figure 2**). To test the specificity of FSEN1, we examined its ability to inhibit NQO1, another CoQ oxidoreductase that has been implicated in ferroptosis^34^. In contrast to FSP1 (**Figure 3A**), FSEN1 had no effect on the CoQ oxidoreductase activity of NQO1 (**Figure 3B**), indicating that FSEN1 does not generally inhibit CoQ oxidoreductases and that FSEN1 exhibits selectivity towards FSP1.

**Figure 3.**
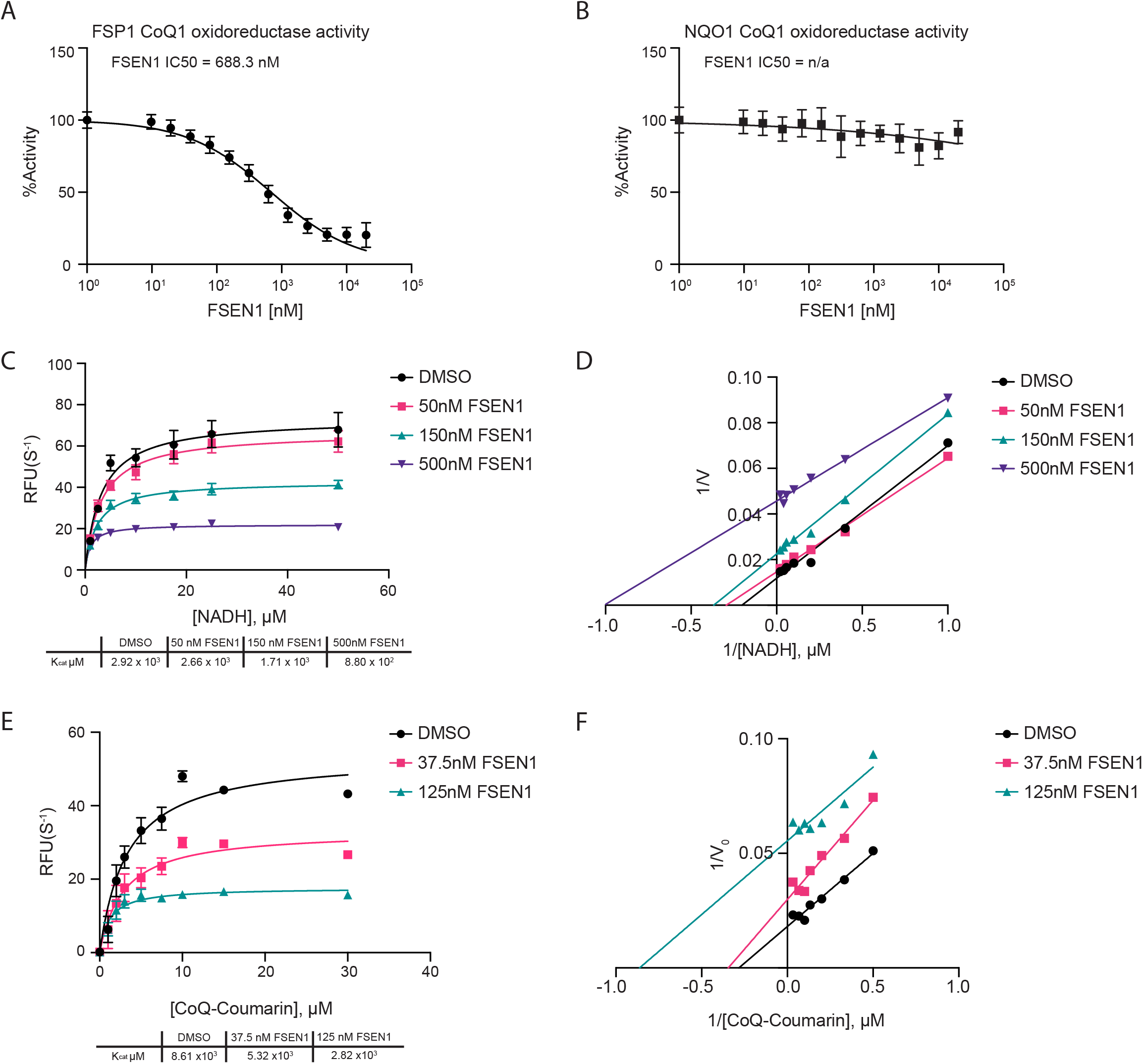
Mechanisms of inhibition of FSP1 by FSEN1. **A,B)** Purified recombinant FSP1 (**A**) and NQO1 (**B**) CoQ1 oxidoreductase activities were measured in the presence of increasing concentrations of FSEN1. Data was normalized to the slopes calculated from DMSO and No Protein controls. IC50 values displayed in nM were calculated from anon-linear regression curve fit. Data are mean ± SEM bars (n=3). **C,D)** Michaelis-Menten and Line Weaver-Burk plots of FSP1 treated with increasing concentrations of NADH in the presence of vehicle or FSEN1. 10 µM CoQ-Coumarin was used as the co-substrate for FSP1, and reduced CoQ-Coumarin fluorescence was measured as a read-out of enzymatic product formation. Error bars represent linear regression standard error of initial rates taken from three biological replicates. Vmax and Km were calculated from the non-linear regression curve fit. **E,F)** Michaelis-Menten and Line Weaver-Burk plots of FSP1 treated with increasing concentrations of CoQ-Coumarin in the presence of vehicle or FSEN1. 200 µM NADH was included as a co-substrate for FSP1, and reduced CoQ-Coumarin fluorescence was measured as the read-out of enzymatic product formation. Error bars represent linear regression standard error of initial rates taken from three biological replicates. Vmax and Km were calculated from the non-linear regression curve fit.

To understand the mechanism of FSP1 inhibition by FSEN1, enzyme kinetics were analyzed using the *in vitro* FSP1 activity assay in the presence of increasing amounts of its substrates, NADH (**Figure 3C,D**) and a fluorescent ubiquinone analogue CoQ-coumarin (**Figure 3E,F**). As FSEN1 concentrations increased, we observed a decrease in Kcat values (**Figure 3C,E**), and the slopes of the Lineweaver-Burke plots were parallel (**Figure 3D,F**). We observed a similar effect of FSEN1 on the Vmax of FSP1 using CoQ1 as a substrate (**Figure S4**), but this assay was unable to resolve the Km due to limitations in its sensitivity. These findings reveal that FSEN1 is an uncompetitive inhibitor of FSP1. Thus, FSP1 requires binding to its substrates NADH and CoQ first in order to be permissive for FSEN1 binding, which then yields the inactive complex.

### FSEN1 triggers ferroptosis in cancer cells by inhibiting FSP1

To characterize the cell death induced by FSEN1 treatment, we further examined its effects on H460^C^ Cas9 lung cancer cells. Treatment with FSEN1 sensitized H460^C^ Cas9 cells to cell death induced by two GPX4 inhibitors, RSL3 and ML162 (**Figure 4A-C**). A checkerboard dose-response matrix of FSEN1 (0-15 µM) and RSL3 (0-15 µM) provided further evidence of synthetic lethality and indicated that the minimal doses that achieve maximal synergy for inducing ferroptosis in H460^C^ Cas9 cells are 0.55 µM FSEN1 together with 0.55 µM RSL3 (**Figure 4B**). Dying cells exhibited characteristic morphologies of ferroptotic cells and the cell death was blocked by ferrostatin-1 (Fer-1) (**Figure 4C**). Consistent with FSEN1 sensitizing these cells to ferroptosis, cell death induced by the co-treatment of FSEN1 and RSL3 was blocked by incubation with known ferroptosis inhibitors, including the radical trapping antioxidants idebenone, Fer-1, and tocopherol, and the iron chelator deferoxamine (DFO) (**Figure 4D, FigureS5D**). In contrast, the apoptosis inhibitor Z-VAD-FMK and necroptosis inhibitor Nec1s had no effect (**Figure 4D, FigureS5D**). Furthermore, treatment with FSEN1 sensitized cells to lipid peroxidation, as measured using the fluorescent lipid peroxidation reporter dye BODIPY C11 (**Figure 4E**). These findings demonstrate that FSEN1 sensitizes H460 lung cancer cells to lipid peroxidation and ferroptosis induced by GPX4 inhibition.

**Figure 4.**
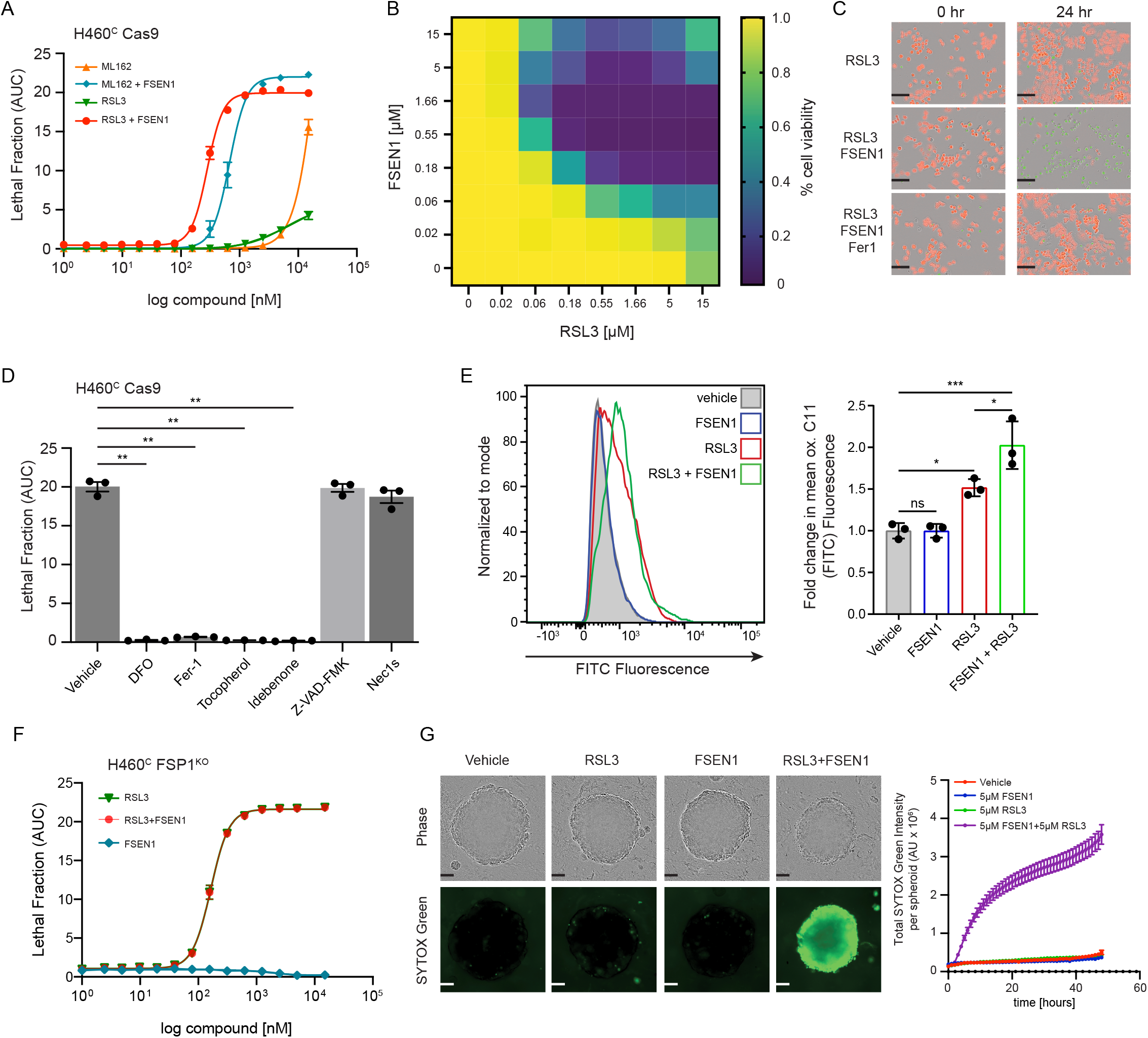
FSEN1 is synthetic lethal with GPX4 inhibitors and sensitizes cancer cells to ferroptosis through inhibition of FSP1. **A)** Dose response of RSL3 and ML162-induced cell death in H460^C^ Cas9 cells co-treated with 1 µM FSEN1 and 2 µM Fer-1 where indicated. Data are mean ± SEM from three biological replicates. **B)** Heat map represents the fraction of viable H460^C^ Cas9 cells co-treated with increasing doses of FSEN1 and RSL3. Data are from three biological replicates. **C)** Representative images of H460^C^ Cas9 cells co-treated with 1.6 µM RSL3, 1 µM FSEN1 and 2 µM Fer-1 as indicated. Scale bar = 200 µm. **D)** Lethal Fraction (AUC) of 5 µM RSL3-induced cell death in H460C Cas9 cells co-treated with 1 µM FSEN1 together with the indicated inhibitors of ferroptosis (Fer-1 [2 µM], DFO [100 µM], idebenone [10 µM], tococopherol [10 µM]), apoptosis (Z-VAD [10 µM]), and necroptosis (Nec1s [1 µM]). Data are mean ± SEM from three biological replicates. **p<0.01 by one-way ANOVA with Dunnett’s multiple comparisons test. **E)** Representative flow cytometry histogram (left panel) and quantification (right panel) of H460^C^ Cas9 cells treated with 200 nM RSL3 and/or 1 µM FSEN1 and labeled with the lipid peroxidation sensor BODIPY 581/591 C11. Green fluorescence intensity was analyzed by flow cytometry with the FITC channel. Data are mean ± SD of three biological replicates. *p<0.05, ***p=0.0003 by one-way ANOVA with Tukey’s multiple comparisons test. **F)** Dose response of RSL3-induced cell death in H460^C^ FSP1^KO^ cells co-treated with 1 µM FSEN1. Data are mean ± SEM from three biological replicates. **G)** H460^C^ spheroids were treated with vehicle, 5 µM FSEN1, and 5 µM RSL3 as indicated. Representative images are shown. Scale bar represents 100 µm. The total intensity of the SYTOX green signal for each spheroid was quantified and the mean ± SEM plotted (n = 20 independent spheroids).

To determine if FSEN1 sensitizes cells to ferroptosis by inhibiting FSP1, we tested the effect of FSEN1 on RSL3-induced ferroptosis in a H460^C^ cell line in which we knocked out FSP1 using CRISPR-Cas9 (H460^C^ FSP1^KO^ cells). H460^C^ FSP1^KO^ cells were greatly sensitized to RSL3-induced cell death (**Figure 4F**). Importantly, although FSEN1 sensitized H460^C^ Cas9 cells to RSL3-induced cell death, treatment of the H460^C^ FSP1^KO^ cells with FSEN1 did not result in any additional sensitization to RSL3-induced cell death (**Figure 4F**). These data indicate that the effects of FSEN1 are due to on target inhibition of FSP1 and not through inhibition of other ferroptosis resistance factors, including other CoQ oxidoreductases implicated in ferroptosis such as NQO1^34^.

High cell densities and cell-cell interactions promote ferroptosis resistance by inducing multiple signaling pathways, including the NF2–YAP^35^ and TAZ-EMP1-NOX4^36^ pathways, underscoring the importance of testing ferroptosis sensitivity in 3-dimensional (3-D) tumor models. Spheroids are 3-D aggregates of cancer cells that more closely reflect key characteristics of solid tumor biology, including cell-cell interactions, hypoxia, drug penetration, and interactions with deposited extracellular matrix^37^. Importantly, similar to our 2-D culture experiments, RSL3 and FSEN1 synergized to trigger cell death in H460 cells grown in the 3-D spheroids (**Figure 4G**). Together, these findings indicate that FSEN1 sensitizes H460 lung cancer cells to ferroptosis in both 2-D and 3-D culture models.

### FSEN1 sensitizes multiple cancer cell lines of different tissue origins to ferroptosis

To examine the role of FSP1 in suppressing ferroptosis in different types of cancer, we measured the impact of FSEN1 on RSL3-induced cell death in a panel of cancer cell lines of various tissue origins, including lung (A549^N^), breast (HCC1143^C^), liver (Huh7^C^), glial (T98G^N^), bone (U 2-OS^N^), connective (HT-1080^N^), lymphoid (RL, SUDHL5), and skin (A375, SKMEL28, 501-MEL) (**Figure 5A,B and Figure S5A,B**). FSEN1 sensitized all cancer cells to RSL3-induced ferroptosis to varying extents, and in all cases the cell death was rescued by co-treatment with Fer-1. These findings indicate that FSEN1 can sensitize multiple cancer cell lines of different origins to ferroptosis induced by GPX4 inhibition. Notably, FSEN1 had a particularly large sensitizing effect on ferroptosis induction in A549 lung cancer cells (**Figure 5A**). A549, and the H460 cells used in our initial screens, are both lung cancer cell lines with KEAP1 mutations, which results in NRF2-dependent upregulation of FSP1^29^. The prominent role of FSP1 in protecting A549 and H460 cells from ferroptosis correlated with high FSP1 protein levels and low GPX4 levels relative to other cancer cell lines (**Figure S5C**). It was also notable that FSEN1 induced a small amount of ferroptosis in A375 melanoma cells in the absence of an RSL3 co-treatment (**Figure 5B**), indicating that A375 cells have a particularly strong dependence on FSP1 for ferroptosis suppression. The amount of sensitization imbued by FSEN1 likely depends on the expression levels of FSP1 and ferroptosis related factors, such as GPX4, DHODH, GCH1, ACSL3, ACSL4, and others.

**Figure 5.**
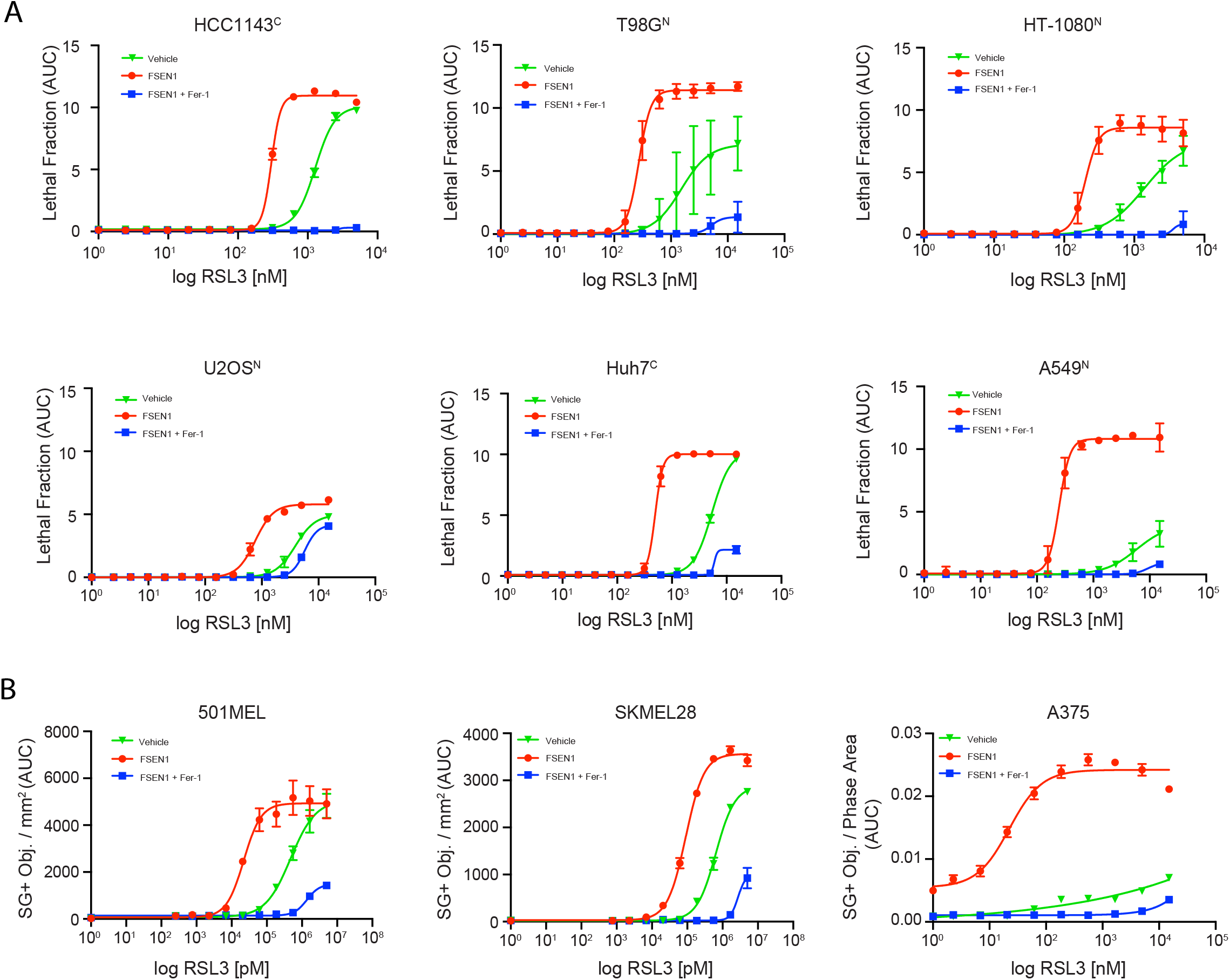
FSEN1 sensitizes cancer cells from different origins to ferroptosis. **A)** Dose response of RSL3-induced cell death in the indicated cancer cell lines treated in the presence and absence of 1 µM FSEN1 and 2 µM Fer-1 as indicated. Data are mean ± SEM from three biological replicates. **B)** Quantification of cell death in melanoma cell lines treated as in (**A**), calculated as SYTOX green positive object per mm^2^. Data are mean ± SEM of two biological replicates.

### FSP1 and dihydroartemisinin treatment synergize to trigger ferroptosis in cancer cells

As ferroptosis suppression is mediated by several different pathways, it is perhaps not surprising that FSP1 inhibition alone is not sufficient to induce ferroptosis in most cancer cell lines. Furthermore, while FSEN1 is synthetically lethal with GPX4 inhibitors such as RSL3 (**Figure 4A**), the *in vivo* efficacy of RSL3 is known to be limited due to low solubility and poor^28^. Therefore, we conducted a synthetic lethal screen to identify compounds that eliminate cancer cells specifically in presence of FSEN1. H460^C^ FSP1^KO^ cells were treated for 24 h with a library of 5,370 compounds that includes 1,200 FDA approved drugs and 4,170 bioactive compounds, and lethal fraction was quantified by time-lapse fluorescence imaging (**Figure 6A,B**). RSL3 was included as a positive control. As expected, RSL3 was cytotoxic for the H460^C^ FSP1^KO^ cells but not the H460^C^ Cas9 cells (**Figure 6B**), validating the ability of our screening approach to detect synthetic lethal relationships.

**Figure 6.**
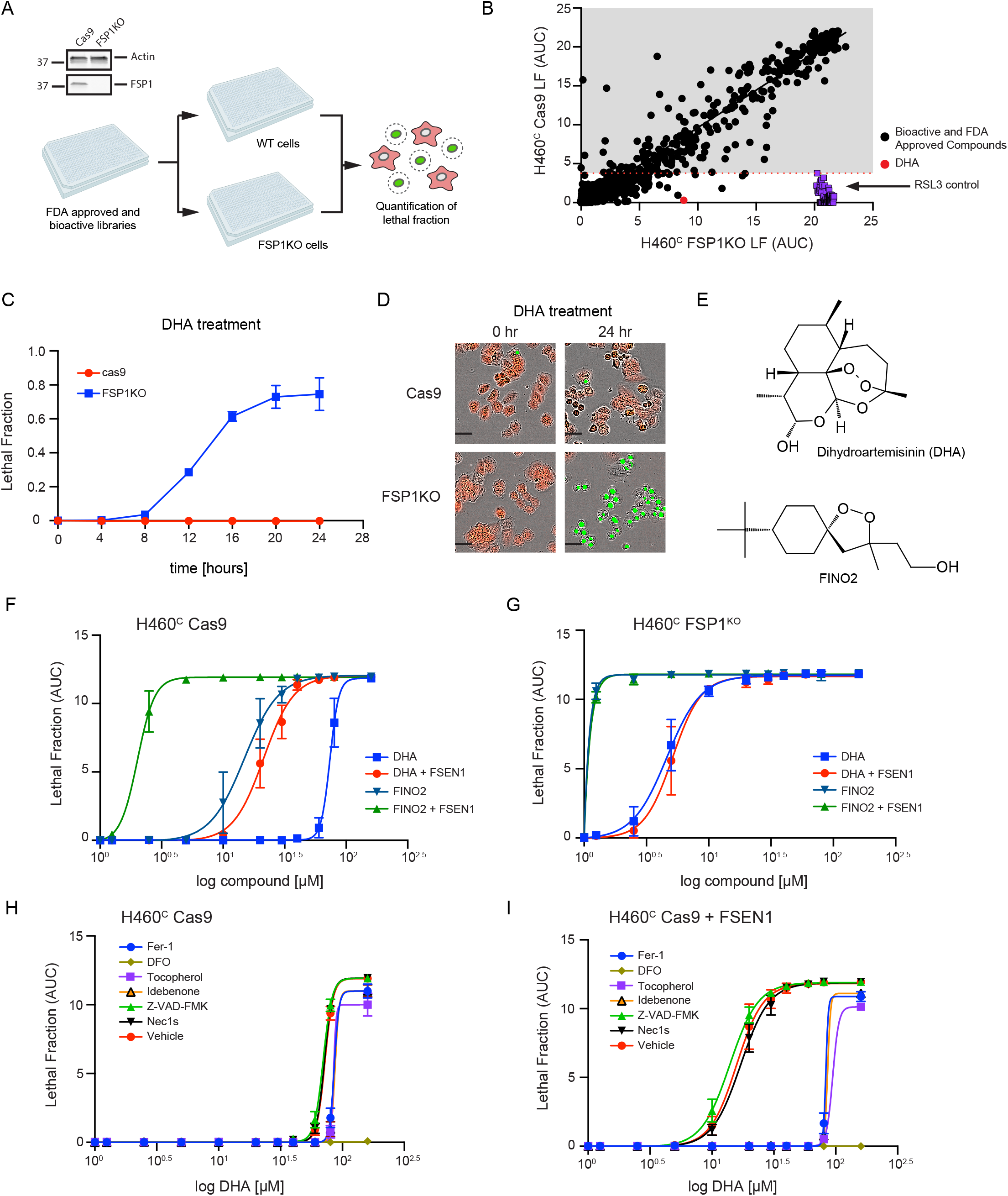
FDA library screens identify DHA as inducer of ferroptosis in FSP1KO cells. **A)** Screen schematic. H460^C^ Cas9 and FSP1^KO^ cells were incubated with 40 µM compounds and lethal fraction was quantified. **B)** Scatter plot of the lethal fraction (LF) for each compound in H460^C^ Cas9 and H460^C^ FSP1^KO^ cells (n=1). Red dotted line represents the cutoff defined by the RSL3 positive control. **C)** Dose response validation of lethal fraction over time comparing H460^C^ Cas9 and FSP1^KO^ cells treated with 40 µM DHA. **D)** Representative IncuCyte images of H460^C^ Cas9 and FSP1^KO^ cells treated with DHA. Scale bar represents 50 µm. **E)** The chemical structures of DHA and FINO2. **F,G)** Dose response of DHA and FINO2-induced cell death in H460^C^ Cas9 (**F**) and FSP1^KO^ (**G**) cells co-treated with vehicle or 1 µM FSEN1. Data are mean ± SEM bars from two biological replicates. **H,I)** Dose response of DHA-induced cell death in H460^C^ cells co-treated with vehicle (**H**) or 1 µM FSEN1 (**I**) together with the indicated inhibitors of ferroptosis (Fer-1 [2 µM], DFO [100 µM], idebenone [10 µM], tococopherol [10 µM]), apoptosis (Z-VAD [10 µM]), and necroptosis (Nec1s [1 µM]). Data are mean ± SEM from two biological replicates.

Several compounds selectively induced cell death in the FSP1^KO^ cells, including DHA (**Figure 6B-D**). DHA is a sesquiterpene lactone compound (**Figure 6E**) that has been widely used as an antimalarial and has also been explored as an anti-cancer therapeutic^38^. The endoperoxide bridge within DHA (**Figure 6E**) is known to react with ferrous iron and stimulate the formation of toxic free radicals^38^. In addition, DHA was recently shown to induce ferroptosis in multiple cancer types^39–42^. Similar to DHA, the ferroptosis inducer FINO2 contains an endoperoxide bridge (**Figure 6E**) that oxidizes iron, increases lipid peroxidation, and induces ferroptosis^43^. FSEN1 treatment and FSP1 KO strongly sensitized H460^c^ cells to cell death induced by both DHA and FINO2 (**Figure 6F,G**). FSEN1 had no additional sensitizing effect in the FSP1^KO^ cells, indicating that the ability of FSEN1 to sensitize cells to DHA and FINO2 induced cell death is due to its on target inhibition of FSP1 (**Figure 6F,G**). DHA and FSEN1 treatment together induced cell death (DHA EC50 = 21.3 µM) that was strongly suppressed by ferroptosis inhibiting radical trapping antioxidants (Fer-1, tocopherol, idebenone) and completely suppressed by DFO (**Figure 6H,I**).

To characterize the synergy between DHA, RSL3, and FSEN1, we performed a checkerboard dose-response matrix of DHA (0-80 µM) and RSL3 (0-15 µM) in the presence of increasing doses of FSEN1 (0, 0.05, 0.5, 5 µM), and quantified synergy potency using a computational zero interaction potency (ZIP) modeling system (**Figure 7A**). The ZIP synergy scoring system defines compound synergy as a value >10, additive effects -10<0<10, and antagonistic effects <0 (**Figure 7A**). The combinations of FSEN1 and RSL3 (ZIP score = 38.28) and FSEN1 and DHA (ZIP score = 26.45) exhibited strong synergy (**Figure 7B,C and Figure S6**). In contrast, DHA and RSL3 showed little synergy (ZIP score = 4.11) (**Figure 7B,C and Figure S6**) and a stronger synergy score was observed with all three compounds (**Figure 7D**). Together, these data indicate that FSP1 inhibition increases cancer cell sensitivity to endoperoxide-containing ferroptosis inducers, including the FDA approved drug DHA. Moreover, these findings demonstrate the potential value of combinatorial therapeutic strategies that combine FSP1 inhibitors with additional ferroptosis inducers to overcome the multitude of cancer ferroptosis defenses.

**Figure 7.**
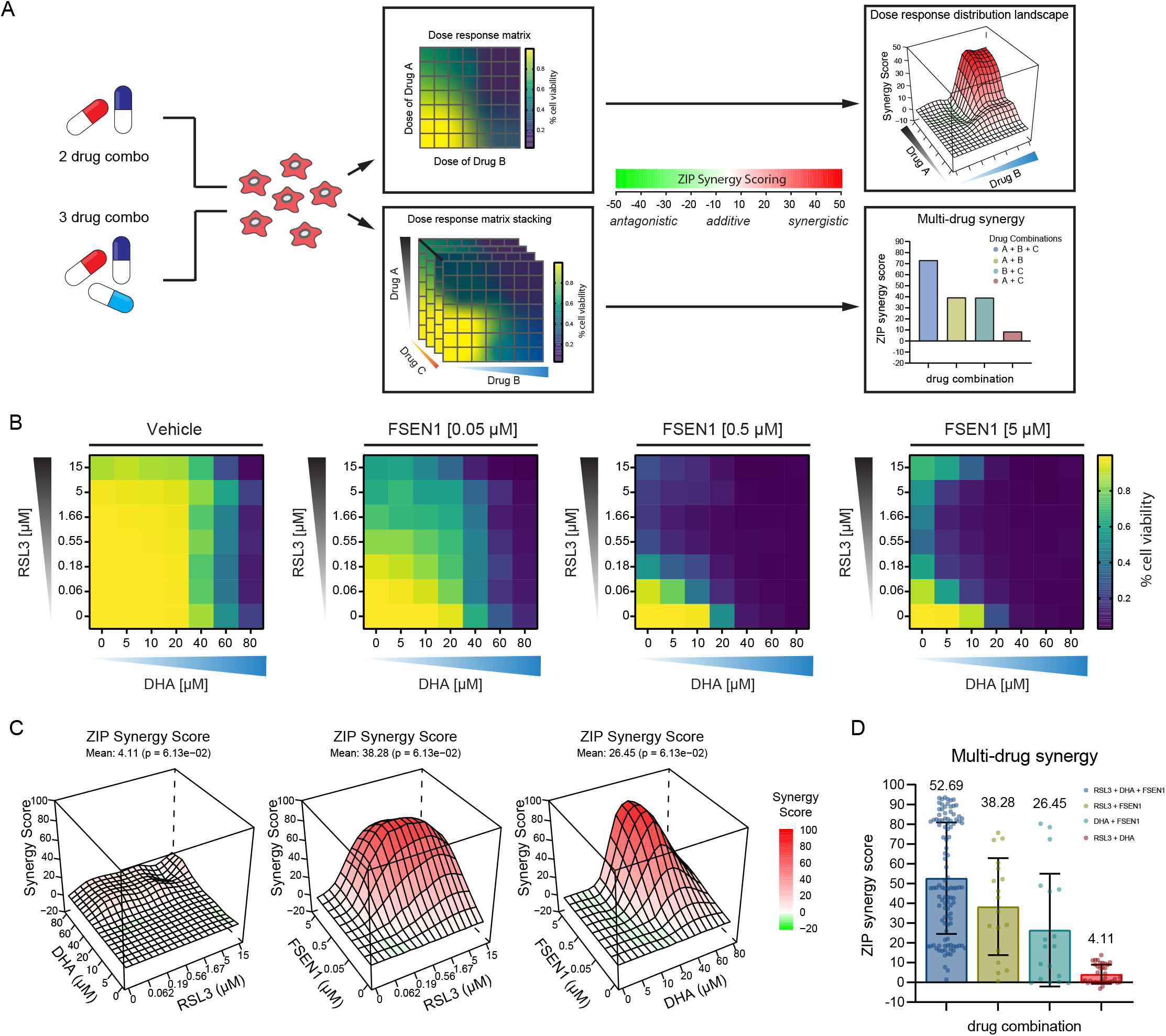
FSEN1 is synergistic with DHA. **A)** Overview of the experimental design and analysis of synergy for two and three drug combinations. **B)** Heat maps of the fraction of viable cells in H460^C^ Cas9 cells co-treated with vehicle and varying doses of FSEN1 (0.05, 0.5, 5 µM) in the presence of increasing concentrations of DHA and RSL3. **C)** 3D-dose response landscape plots illustrating the distribution and depth of synergy between the different drug combinations and doses tested in H460^C^ Cas9 cells with the corresponding average ZIP synergy scores and p-values. **D)** Multi-drug synergy bar plots of H460^C^ Cas9 cells illustrating mean ZIP synergy scores for all possible drug combinations tested. Data are mean ± SEM from three biological replicates. P-values represents the significance of the difference between the estimated average synergy score over the whole dose-response matrix and 0% inhibition under the null hypothesis of non-response.

## DISCUSSION

Induction of ferroptosis has emerged as a promising strategy to treat therapy-resistant cancer cells^14^. Despite the discovery of multiple cellular ferroptosis defense systems^2,14^, current ferroptosis inducers are mostly limited to the GSH-GPX4 pathway, impeding the study and assessment of other ferroptosis regulators as therapeutic targets. Here, we describe the discovery and characterization of structurally distinct small molecule FSP1 inhibitors that directly inhibit FSP1 activity and sensitize cancer cells to ferroptosis. The most potent of these new FSP1 inhibitors, FSEN1, is an uncompetitive FSP1 inhibitor that exhibits synthetic lethality with GPX4 inhibitors. We further find that FSEN1 is synthetic lethal with endoperoxide-containing ferroptosis inducers, including FINO2 and the FDA approved compound DHA.

Given that FSP1 expression correlates with resistance to GPX4 inhibitors across many cancer cell lines^4,5^, it is likely that FSEN1 will be useful for sensitizing many cancer types to ferroptosis inducers that target the GSH-GPX4 axis. Indeed, FSEN1 sensitized several cancer cell lines of different tissue origins bearing different oncogenic mutations. The variable amount of sensitization FSEN1 confers on distinct cancer cell lines indicates that some cancers are more reliant on FSP1 for ferroptosis suppression than others, likely reflecting the contribution of other ferroptosis defense pathways or differences in phospholipid composition and ROS production. Strong sensitization was observed in H460 and A549 cancer cells, which are both lung cancer cell lines with KEAP1 mutations. KEAP1 regulates a canonical pathway that mediates the ubiquitin-dependent proteasomal clearance of NRF2, a master transcriptional factor that governs expression of antioxidant factors, including FSP1^29^. We find that these cell lines have high FSP1 and low GPX4 protein levels, suggesting a means to stratify cancers that will respond more strongly than others to FSP1 inhibition based upon FSP1 and GPX4 levels. A similar increased effectiveness of DHODH inhibition on ferroptosis induction in cancer cells with low GPX4 was previously observed^6^.

FSP1 inhibition alone was not sufficient to trigger ferroptosis in most of the cell types under the culture conditions that we examined here, except for the A375 melanoma cells. It is worth noting that FSP1 KO dramatically slowed tumor growth in an *in vivo* xenograft model of KEAP1 deficient lung cancer^29^, indicating that FSP1 inhibition is sufficient to induce ferroptosis under some *in vivo* conditions. This may reflect the unique contribution of the in vivo tumor microenvironment and higher levels of PUFAs and ROS. Importantly, the observation that FSP1 KO impairs tumor growth *in vivo* raises the possibility that small molecule FSP1 inhibition may be effective as a monotherapy for specific cancers. The result also highlights the importance of future studies to explore FSP1 inhibition in *in vivo* cancer models.

Our findings are consistent with the utility of FSP1 inhibition in combinatorial therapeutic regimes together with inhibitors of the GSH-GPX4 pathway. It is likely that FSP1 inhibition will sensitize cancer cells to standard of care treatments that trigger ROS generation and ferroptosis such as radiotherapy^44,45^, photodynamic therapy^46,47^, and immunotherapy^48^. Indeed, FSP1 KO and iFSP1 treatment sensitize cancer cells to radiotherapy^29^. Along these lines, we discover that FSP1 inhibition synergizes with DHA to induce ferroptosis. DHA is a commonly used antimalarial that has been explored as an anticancer drug^38^, and several recent studies indicate that DHA is capable of triggering ferroptosis types^39–42^. The endoperoxide bridge within DHA reacts with and oxidizes iron, leading to the production of reactive oxygen species and lipid peroxides^38^. In H460 lung cancer cells, DHA induced minimal amounts of ferroptosis on its own, but its ability to induce ferroptosis was greatly enhanced by co-treatment with FSEN1. It remains possible that DHA operates through multiple mechanisms, oxidizing iron and triggering lipid peroxide formation and indirectly inhibiting GPX4 similar to what was reported for the endoperoxide ferroptosis inducer FINO2^43^. Our findings highlight the potential of endoperoxide containing compounds (e.g., FINO2^43^, FINO3^49^, DHA) as therapeutic agents to induce ferroptosis by increasing iron oxidation, lipid peroxidation, and potentially GPX4 inhibition.

In summary, our findings identify a set of structurally distinct FSP1 inhibitors that are effective in overcoming ferroptosis resistance in several cancer cell lines. Future studies will be necessary to determine whether these FSP1 inhibitors are effective in preclinical mouse models of cancer, as single agent therapeutics under certain conditions, and in sensitizing cancer cells to other treatments that increase ROS and lipid peroxidation, including ferroptosis induced by photodynamic therapy, radiation, and immunotherapy.

## ACKNOWLEDGEMENTS

This research was supported by grants from the National Institutes of Health (R01GM112948 to J.A.O. and 1R01GM122923 to S.J.D.), American Cancer Society (Research Scholar Award RSG-19-192-01 to J.A.O. and RSG-21-017-01-CCG to S.J.D.), and Melanoma Research Alliance (Award 620458 to J.A.O.). J.A.O. is a Chan Zuckerberg Biohub investigator and Miller Institute Professor. J.M.H. was supported by a University of California Cancer Research Coordinating Committee (CRCC) predoctoral fellowship. C.D. was supported by a Cayman Biomedical Research Institute (CABRI) Undergraduate Fellowship.

## AUTHOR CONTRIBUTIONS

J.M.H., K.B., J.A.O., and J.S. conceived of the project and designed the experiments. J.M.H. and J.A.O. wrote the manuscript. All authors read, edited, and contributed to the manuscript. J.M.H. performed the majority of the experiments. J.M.H., K.B., and E.W. performed the small molecule screens and analyzed the data. Z.L. performed spheroid assays. J.M.H. and M.R. performed lipid peroxidation assays. I.L.O. assisted with the analysis of melanoma lines. J.M.H. and M.A.R. performed the BODIPY C11 experiments. S.J.D., K.K.D., and M.L. provided critical reagents and guidance. C.D., J.M.H., and K.B. purified proteins and performed *in vitro* analyses of FSP1 activity.

## AUTHOR INFORMATION

Correspondence and requests for materials should be addressed to J.A.O. and J.S.

## METHODS

### Cell lines and culture conditions

A549^N^, HCC1143^C^, NCI-H460^C^, NCI-H460^C^ GPX4^KO^ and FSP1^KO^ lines were cultured in RPMI1640 with l-glutamine (corning). HT-1080^N^, U-2OS^N^, T98G^N^, Huh7^C^, and A375 cells were cultured in DMEM with l-glutamine and without sodium pyruvate (corning). 501MEL and SKMEL28 cells were cultured in DMEM high glucose with l-glutamine and without sodium pyruvate (corning). Cells (HT-1080^N^, U-2OS^N^, T98G^N^) were generated through stable expression of nuclear-localized mKate2 (denoted by an additional superscript ‘N’). All media were supplemented with 10% fetal bovine serum (FBS, Thermo Fisher Scientific and Gemini Bio Products), and all cell lines were grown at 37□°C with 5% CO_2_. All cell lines were tested for mycoplasma and were not authenticated.

### Generation of cell lines

NCI-H460^C^ FSP1^KO^ lines were generated by infection with lentiCRISPR v2-Blast (Addgene plasmid no. 83489) virus, and NCI-H460^C^ GPX4^KO^ lines were generated by infection with lentiCRISPR v2-Hygro (Addgene plasmid no. 98291) virus, as previously described^2^. Cells expressing sgSAFE-mCherry (denoted by an additional superscript ‘C’) were generated from the respected parental cells via transduction with the lentiviral sgRNA expression vector with mCherry pMCB320 (Addgene plasmid no. 89359) virus, which directs the expression of cytosolic sgSAFE-mCherry. Polyclonal sgSAFE-mCherry expressing cells were selected for using puromycin or hygromycin respectively. H460C polyclonal pools were selected puromycin and further enriched using FACS (UC Berkeley shared FACS Facility).

### Plasmids

For protein expression, FSP1(WT), lacking the ATG start codon were inserted into the pET-His6-TEV vector (Addgene plasmid no. 29653), as previously described^2^. For protein expression of NQO1, NQO1(WT) lacking the ATG start codon was generated by PCR amplification of NQO1 from an NQO1-GFP pcDNA5/FRT/TO plasmid generated in our previous study (2). The NQO1(WT) amplicon was inserted into the pET-His6-TEV vector (Addgene plasmid no. 29653), C-terminal to the His6-TEV tag using restriction enzyme-independent fragment insertion by polymerase incomplete primer extension.

### Chemicals and reagents

Reagents used in this study include: RSL3 (Cayman Chemical), Fer-1 (Cayman Chemical), idebenone (Cayman Chemical), DFO (Cayman Chemical), ML162 (Cayman Chemical), ZVAD(OMe)-FMK (Cayman Chemical), necrostatin-1 (Cayman Chemical), DHA (Selleck Chemical), puromycin (Thermo Fisher Scientific), SYTOX Green Dead Cell Stain (Thermo Fisher Scientific), polybrene (Sigma-Aldrich), coenzyme Q_1_ (Sigma-Aldrich), resazurin (Thermo Fisher Scientific) and NADH (Sigma-Aldrich). A 100,000-member compound Diverse Library and a 15,000 compound Antibacterial Library were obtained from ChemDiv. The 1,200 compound FDA approved, and 4,170 compounds Bioactive libraries were obtained from TargetMol. Compounds were stamped into 384 well Non-Binding Surface (NBS) plates (Corning) using a Cybio Well Vario liquid handler (Analytik-Jena, Germany).

### Small molecule screen for FSP1 inhibitors / Cell death analysis

For small molecule drug screen, 0.5 μL of compounds were stamped into 384 well NBS plates (Corning) in dose response diluted 2-fold with a high of 2 mM. 60 mL of purified His tagged FSP1(WT) protein at 50 nM was prepared and 12.5 μL of this protein solution was aliquoted into 384 well NBS-plates (Corning) containing compound and allowed to incubate for 30 min at room temperature. After 30 min incubation 12.5 μL of reaction buffer (1 mM NADH, 800 μM CoQ1) was added into the 384 well NBS-plates containing protein and compounds for a final concentration of 25 nM His tagged FSP1(WT) protein, 500 μM NADH (Sigma-Aldrich) and 400 μM CoQ1 (Sigma-Aldrich). Each well contained a 25 μL mixture and 40 μM compound in the primary screen. All aliquots for in-vitro drug screen were completed with an Analytik-Jena Cybio Well Vario liquid handler. 2 mM compounds were stored in 100% DMSO in 384 well plates. Wells were homogenized with a Bioshake 3000 ELM orbital shaker at 2,400 rpm for 45 s and condensates were allowed to settle for 30 min before scanning. Measurements were taken at 355 nm with a EnVision 2104 multilabel plate reader (PerkinElmer). No protein data control was used for background subtraction prior to upload to CCDvault for normalization to DMSO vehicle control wells. Compounds with confirmed data that have normalized absorbance values of less than 0.164 were chosen for dose response. Wells that exhibit 3 standard deviations from the untreated sample and a no protein control were selected for dose-response screening. The same His-tagged FSP1(WT) protein were tested against candidate drugs using a 10-point serial dilution starting at 40 μM using the same procedure.

For cell-based screen, cells were seeded in triplicate at a density of 500-750 cells per 50 µL per well in black 384-well plates (NUNC) and (Corning) 24 hr before start of imaging. After 24 hr, an additional 50 µL of drug infused medium containing 30 nM SYTOX Green Dead Cell Stain was carefully placed into the wells on top of existing medium. The plates were immediately transferred to an IncuCyte S3 imaging system (Essen Bioscience) enclosed in an incubator set to 37□°C and 5% CO_2_. One image per well were captured in the phase, green, and red channels every 1.5 or 3 hr over a 24 hr period, and the ratio of SYTOX Green-positive objects (dead cells) to SYTOX Green-positive plus Sg-SAFE mCherry-positive objects (total cells) was quantified using S3 image analysis software (Essen Bioscience). For each treatment condition, the SYTOX-to-‘SYTOX+mCherry’-object ratio was plotted against the 24 hr imaging interval, the Area Under the Curve (AUC) was calculated and the average AUC was plotted using Prism (GraphPad). To calculate the half-maximal effective concentration (EC50) values, the AUC curve was fit to a variable slope function comparing response to drug concentration.

### Lipid peroxidation assay

Cells seeded in a 6-well plate were treated with 200 nM RSL3 for 5 hr and washed once with DPBS containing calcium and magnesium. Cells then were incubated in DPBS containing 5 μM BODIPY 581/591 C11 (Invitrogen, #D3861) at 37 °C for 10 min and washed 3x with DPBS without calcium or magnesium. Cells were detached from the plate with trypsin, and green fluorescence was analyzed by flow cytometry (>10,000 cells) on a BD LSRFortessa. Data were analyzed using FlowJo.

### Spheroid / 3D cell culture

2500 NCI-H460 cells with 100 µL full serum RPMI media were seeded in 96-well Black/Clear Round Bottom Ultra-Low Attachment Spheroid Microplate (Corning, #4515). Cells were incubated at 37°C for 30min before another 100µL RPMI media containing 2% Matrigel (Corning #354234) was added. Plates were centrifuged at 750x g for 15min and grew at 37 °C for 2 days. For FSEN1 and RSL3 treatment, 100 µL RPMI media was slowly removed without disturbing the spheroid. Another 100 µL RPMI media containing 1% Matrigel, 60 nM SYTOX Green dye, 10 µM FSEN1 or 10 µM RSL3 or both was added back into each well. Spheroids were imaged in IncuCyte S3 with a 10x objective.

### FDA and Bioactive Library Screen

For FDA and Bioactive Library Screen experiments, cells were seeded at a density of 750 cells per 25 uL per well in black 384-well plates (NUNC) and (Corning) 24 hr before start of imaging. After 24 hr, an additional 25 uL of drug infused medium containing 30 nM SYTOX Green Dead Cell Stain was carefully placed into the wells on top of existing medium. The plates were immediately transferred to an IncuCyte S3 imaging system (Essen Bioscience) enclosed in an incubator set to 37□°C and 5% CO_2_. One image per well were captured in the phase, green, and red channels every 4 hr over a 24 hr period, and the ratio of SYTOX Green-positive objects (dead cells) to SYTOX Green-positive plus SgSAFE mCherry-positive objects (total cells) was quantified using S3 image analysis software (Essen Bioscience). For each treatment condition, the SYTOX-to-‘SYTOX+mCherry’-object ratio was plotted against the 24 hr imaging interval, the Area Under the Curve (AUC) was calculated, and the average AUC was plotted as a function of drug concentration (for example, RSL3) using Prism (GraphPad). To calculate the half-maximal effective concentration (EC50) values, the AUC curve was fit to a variable slope function comparing response to drug concentration.

### Synergy experiments

For cell-based synergy experiments, cells were seeded at a density of 750 cells per 25 µL per well in four black 384-well plates (Corning) 24 hr before start of imaging. After 24 hr, an additional 25 µL of drug infused medium containing 30 nM SYTOX Green Dead Cell Stain was carefully placed into the wells on top of existing medium. The plates were immediately transferred to an IncuCyte S3 imaging system (Essen Bioscience) enclosed in an incubator set to 37□°C and 5% CO_2_. One image per well were captured in the phase, green, and red channels every 4 hr over a 24 hr period, and the ratio of SYTOX Green-positive objects (dead cells) to SYTOX Green-positive plus Sg-SAFE mCherry-positive objects (total cells) was quantified using S3 image analysis software (Essen Bioscience). For each treatment condition, the SYTOX-to-‘SYTOX+mCherry’-object ratio was plotted against the 24 hr imaging interval, the Area Under the Curve (AUC) was calculated, and the average AUC was plotted as a function of drug concentration (for example, RSL3) using Prism (GraphPad). To calculate the half-maximal effective concentration (EC50) values, the AUC curve was fit to a variable slope function comparing response to drug concentration.

### Compound preparation for synergy experiments

Vehicle, RSL3 and DHA combinations were stamped in grid format 1:1 at 4x concentrations. The compounds were then diluted with SYTOX green infused media containing either vehicle, 0.10 µM, 1 µM, or 10 µM FSEN1 to the appropriate 2x concentration (PlateOne, #1884-2410). Compounds were then homogenized with a Bioshake 3000 ELM orbital shaker at 2,400 rpm for 45 sec prior to treatment.

### Protein purification

Expression vectors were transformed into LOBSTR-BL21 (DE3) competent cells (Kerafast) and LB broth was inoculated for overnight growth at 37°C while shaking at 280 rpm. The following day, the cultures were diluted 1:100 into 1L LB broth and allowed to grow at 37°C while shaking at 280 rpm until the cultures reached an optical density at 600 nm (OD_600_) of 0.6, measured by NanoDrop One (Thermo Scientific Cat: 13-400-518). The cultures were then induced with 0.7 mM isopropyl β-D-1-thiogalactopyranoside (IPTG) and grown at 30°C while shaking at 280 rpm overnight. Cultures were then centrifuged for 45 minutes at 4°C and 4500 rpm to pellet. 1L Bacterial pellets were resuspended in 25 mL lysis buffer (50 mM KH_2_PO_4_, 300 mM KCl, 10% glycerol, 30 mM imidazole, 1mg/mL lysozyme, and 1mM PMSF, pH 8.0) and incubated at 37°C while shaking at 280 rpm for 10 minutes. The resuspended cells were lysed by passing through a microfluidizer (Microfluidics model: LM10) 3× at 15,000 PSI and collected in an ice-bath-chilled beaker. Lysate was then ultracentrifuged for 30 minutes at 4°C and 50,000 × g. All subsequent purification steps were carried out at 4°C. The supernatant was passed through an Econo-Pac disposable chromatography column (Bio-Rad Cat: 732-1010) packed and equilibrated with 1mL HisPur Ni-NTA agarose resin (Thermo Scientific Cat: 88221) and washed 4× with 2mL EQ buffer (50 mM KH_2_PO_4_, 300 mM KCl, 10% glycerol, and 30 mM imidazole, pH 8.0). Bound proteins were eluted from Ni-NTA resin bed by gravity flowing 5mL elution buffer (50mM KH2PO4, 300mM KCl, 10% glycerol, 250mM imidazole, pH 8.0) through the column. Using an Amicon Ultra-15 centrifugal filter (Sigma-Aldrich SKU: UFC910008), the eluted proteins were buffer exchanged into SEC buffer (50mM HEPES, 100mM KCl, 2mM DTT, pH 8.0), spiked with 15% glycerol and 10mM DTT and further concentrated to 3mL total volume. Protein sample was then further purified by gel filtration through a HiLoad 16/600 Superdex 75 Pg size exclusion chromatography column (Sigma-Aldrich SKU: GE28-9893-33) in SEC buffer using a GE Akta Pure FPLC. Fractions were collected in 1.6mL aliquots and combined based on purity, visualized by Coomassie staining after separation in SDS-page. Combined fractions were then concentrated to 2mg/mL and snap-frozen in 50μL aliquots using liquid nitrogen. Protein concentration was determined using a Pierce bicinchoninic acid (BCA) protein assay kit (Thermo Scientific Cat: PI23227).

### FSP1 Kinetics (NADH)

To measure FSP1 kinetics, FSEN1 was dissolved and diluted in DMSO, and recombinant purified FSP1 was diluted in PBS. 0.5μL FSEN1 and 12.5μL FSP1 were mixed and incubated for 30 minutes in a NBS polystyrene 384-well plate (Corning Ref: 3640) at RT. Vehicle control wells included 0.5μL DMSO and were absent of any FSEN1. NADH (Millipore Cat: 481913) was then dissolved and diluted in PBS and added to the plate wells. CoQ-Coumarin (Cayman Item: 29554) was dissolved in DMSO, diluted in PBS, and added to the plate wells to start the reaction. Final well volume was 25μL, and final concentrations were 25nM FSP1, 10μM CoQ-Coumarin, 1-50μM NADH, and 50-500nM FSEN1. Following addition of CoQ-Coumarin, the well-volumes were mixed by an orbital shaker for 20 seconds at an amplitude of 5mm and 120rpm. Reduced CoQ-Coumarin fluorescence (Ex: 405nm, Em: 475nm) was measured as read-out of enzymatic product formation on a kinetic cycle with an interval time of 20 seconds at RT. All data was acquired using a Tecan infinite M1000. Raw data from 3 biological replicates were then plotted and initial slopes were calculated using a linear regression in prism (GraphPad).

### FSP1 Kinetics (CoQ-Coumarin)

To measure FSP1 kinetics, FSEN1 was dissolved and diluted in DMSO, and recombinant purified FSP1 was diluted in PBS. 0.5μL FSEN1 and 12.5μL FSP1 were mixed and incubated for 30 minutes in a NBS polystyrene 384-well plate (Corning Ref: 3640) at RT. Vehicle control wells included 0.5μL DMSO and were absent of any FSEN1. NADH (Millipore Cat: 481913) was dissolved and diluted in PBS, and CoQ-Coumarin (Cayman Item: 29554) was dissolved in DMSO and diluted in PBS. Diluted NADH and CoQ-Coumarin were combined into a 2X reaction mix and added to the plate wells to start the reaction. Final well volume was 25μL, and final concentrations were 6.25nM FSP1, 1-30μM CoQ-Coumarin, 200μM NADH, and 12.5-125nM FSEN1. Reduced CoQ-Coumarin fluorescence (Ex: 405nm, Em: 475nm) was measured as read-out of enzymatic product formation on a kinetic cycle with an interval time of 15 seconds at RT. All data was acquired using a Tecan Spark. Raw data from 3 biological replicates were then plotted and initial slopes were calculated using a linear regression in prism (GraphPad).

### FSP1 Kinetics (CoQ1)

To measure FSP1 kinetics, FSEN1 was dissolved and diluted in DMSO, and recombinant purified FSP1 was diluted in PBS. 0.5μL FSEN1 and 12.5μL FSP1 were mixed and incubated for 30 minutes in in a NBS polystyrene 384-well plate (Corning Ref: 3640) at RT. Vehicle control wells included 0.5μL DMSO and were absent of any FSEN1. NADH (Millipore Cat: 481913) was dissolved and diluted in PBS, and CoQ1 (Sigma-Aldrich SKU: C7956) was dissolved in DMSO and diluted in PBS. Diluted NADH and CoQ1 were combined into a 2X reaction mix and added to the plate wells to start the reaction. Final well volume was 25μL, and final concentrations were 25nM FSP1, 200-500μM CoQ1, 500μM NADH, and 0.05-1μM FSEN1. NADH Absorbance (355nm) was measured as an inverse read-out of enzymatic product formation on a kinetic cycle with an interval time of 3 minutes at RT. All data was acquired using a Tecan infinite M1000. Raw data from 2 biological replicates were then plotted and initial slopes were calculated using a linear regression in prism (GraphPad).

### FSP1 and NQO1 % Activity Curves

To measure in vitro activity of FSP1 and NQO1 for IC50 calculation, recombinant purified FSP1 and NQO1 were diluted in PBS. FSEN1 was dissolved and diluted in DMSO, and 0.5 μL FSEN1 and 12.5 μL FSP1/NQO1 were mixed on an orbital shaker for 45 seconds at 450 rpm and incubated for 30 minutes in in a NBS polystyrene 384-well plate (Corning Ref: 3640) at RT. NADH (Millipore Cat: 481913) was dissolved and diluted in PBS, and CoQ1 (Sigma-Aldrich SKU: C7956) was dissolved in DMSO and diluted in PBS. Diluted NADH and CoQ1 were combined into a 2X reaction mix and added to the plate wells to start the reaction. Final well volume was 25 μL, and final concentrations were 12.5n M FSP1, 400 μM CoQ1, 500 μM NADH, and 0.01-20 μM FSEN1. NADH Absorbance (355 nm) was measured as an inverse read-out of enzymatic product formation on a kinetic cycle with an interval time of 2.5 minutes at RT. All data was acquired using a Tecan infinite M1000. Raw data from 3 biological replicates were then plotted and initial slopes were calculated using a linear regression in prism (GraphPad), and all rates were normalized to vehicle (0 nM FSEN1) and No Protein controls where the highest and lowest slope values were used as 0 and 100%. Normalized values were then plotted as a function of FSEN1 concentration in log scale, and prism (GraphPad) was used to perform a non-linear regression curve fit for log (inhibitor) vs. normalized (variable) slopes. IC50 values of FSEN1 for FSP1 and NQO1 were obtained from this non-linear regression.

### Western blotting

Cells were washed two times with PBS prior to lysis in 1% SDS. Samples were then sonicated for 30 sec and incubated for 5 min at 100□°C. Protein concentrations were determined using the bicinchoninic acid (BCA) protein assay (Thermo Fisher Scientific), and equal amounts of protein by weight were combined with 1× Laemmli buffer, separated on 4–20% polyacrylamide gradient gels (Bio-Rad Laboratories) and transferred onto nitrocellulose membranes (Bio-Rad Laboratories). Membranes were washed in PBS with 0.1% Tween-20 (PBST) and blocked in PBST containing 5% (w/v) dried milk or 5% (w/v) bovine serum albumin (BSA) for 30 min. Membranes were incubated for 24 hr in PBST containing 5% BSA (Akron Biosciences) and primary antibodies. After washing with PBST, membranes were incubated at room temperature for 30 min in 5% BSA and PBST containing fluorescent secondary antibodies. Immunoblots were imaged on a LI-COR imager (LI-COR Biosciences).

The following blotting reagents and antibodies were used: anti-AMID (Santa Cruz Biotechnology), anti-β-actin (Santa Cruz Biotechnology), anti-GPX4 (Abcam), anti-rabbit IRDye800 conjugated secondary (LI-COR Biosciences) and anti-mouse Alexa Fluor 680 conjugated secondary (Invitrogen).

## SUPPLEMENTAL FIGURE LEGENDS

**Figure S1.**
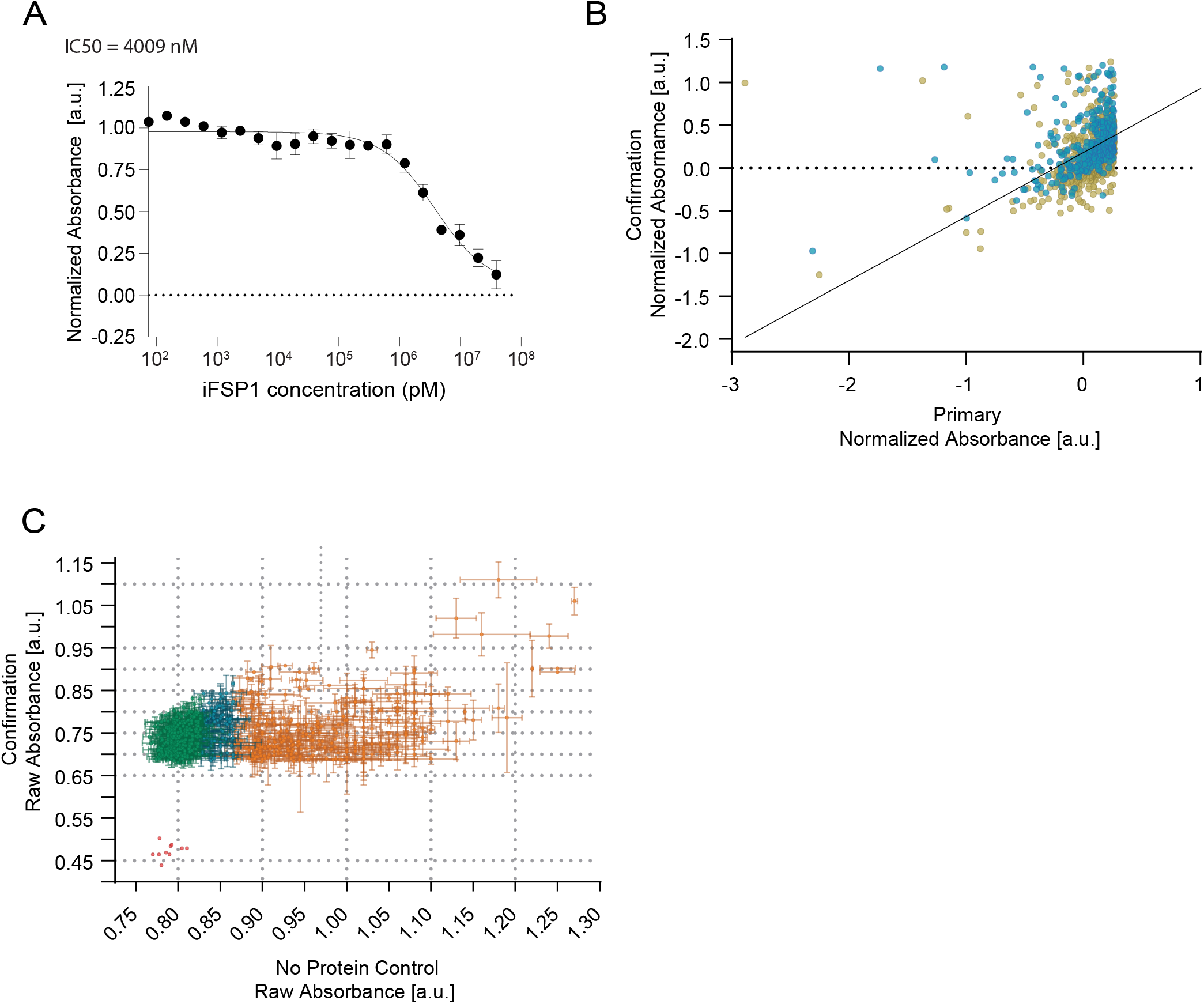
FSP1 *in vitro* activity assay and screen controls. **A)** Dose response analyses were performed for iFSP1 using the *in vitro* absorbance-based assay of FSP1 activity. Data are mean ± SEM bars from two independent experiments. **B)** 1,120 candidate inhibitors were re-tested in duplicate. The confirmation screen is plotted against the primary screen data. 743 of the 1,120 compounds were classified as confirmed “Hits”. **C)** The 743 confirmed compounds (panel B) were tested in assays lacking FSP1 (i.e., no protein control). 323 compounds (green) had activity less than 5% above the DMSO average (red dots) and were selected for further validation. Potential false positives (blue) and false positives (orange) were removed from further analysis.

**Figure S2.**
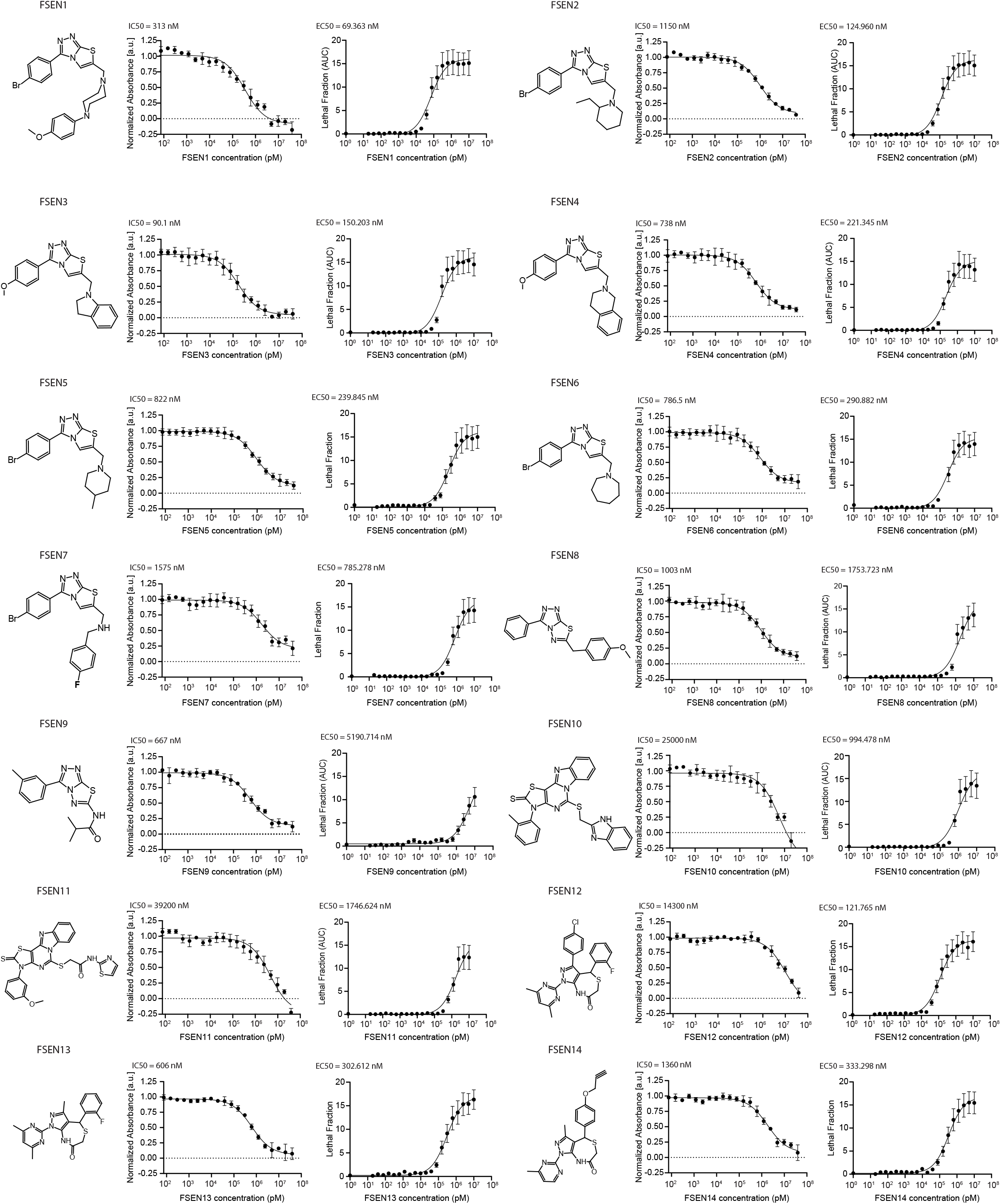
Analysis of FSEN1-14 inhibition of FSP1. Dose response analyses were performed for FSEN1-14 using the *in vitro* absorbance-based assay of FSP1 activity and the H460^C^ GPX4^KO^ cell-based assay of ferroptosis to determine the compound’s IC50 and EC50, respectively. IC50 values were calculated from end-point assays for inhibition of FSP1 activity *in vitro* (mean ± SEM from two independent experiments) and their EC50 values were calculated from lethal fraction (AUC) quantification in H460^C^ GPX4KO cells (data are mean ± SEM from three biological replicates). The IC50 and EC50 values calculated from these experiments are also listed in **Figure 2**.

**Figure S3.**
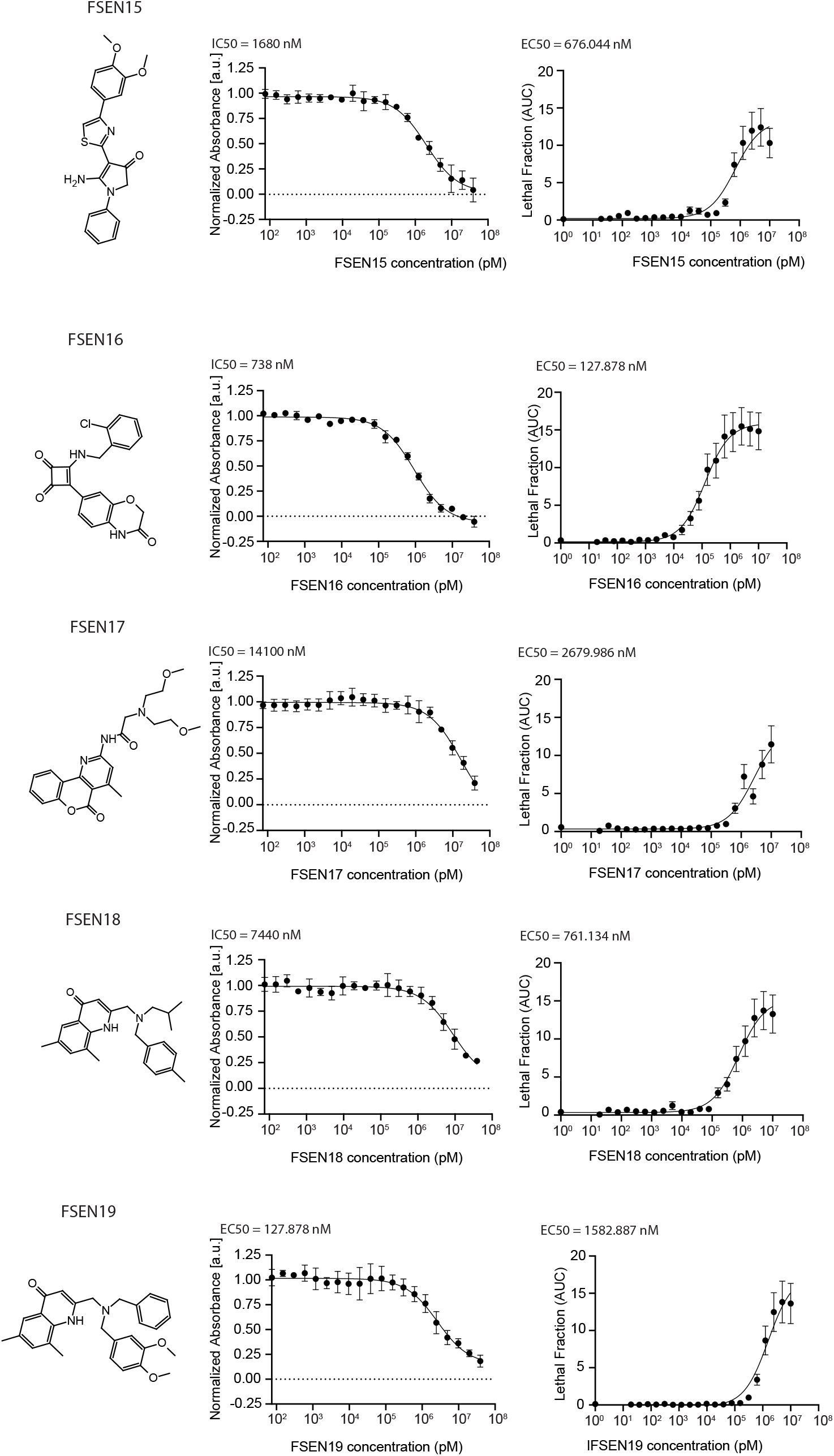
Analysis of FSEN15-18 inhibition of FSP1. Dose response analyses were performed for FSEN15-19 using the *in vitro* absorbance-based assay of FSP1 activity and the H460^C^ GPX4^KO^ cell-based assay of ferroptosis to determine the compound’s IC50 and EC50, respectively. IC50 values were calculated from end-point assays for inhibition of FSP1 activity *in vitro* (mean ± SEM from two independent experiments) and their EC50 values were calculated from lethal fraction (AUC) quantification in H460^C^ GPX4KO cells (data are mean ± SEM from three biological replicates). The IC50 and EC50 values calculated from these experiments are also listed in **Figure 2**.

**Figure S4.**
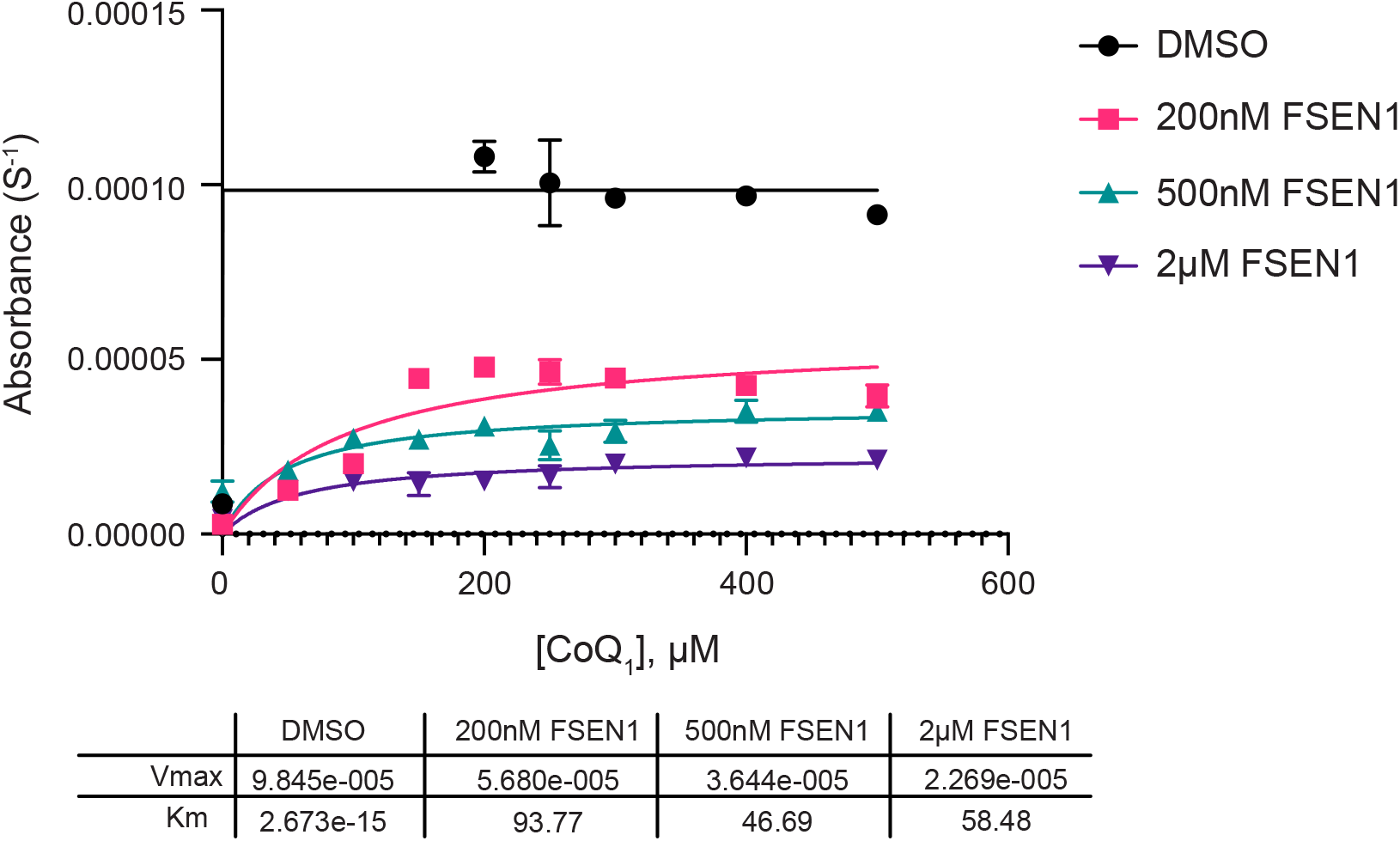
FSEN1 FSP1 Kinetics (CoQ1) Michaelis-Menten Curve of Purified recombinant FSP1 treated with increasing concentrations of CoQ1 in the presence of vehicle or FSEN1 (0.05, 0.15, 1 uM). All NADH concentrations included 400 μM CoQ1 as co-substrate for FSP1, and NADH absorbance (355 nm) was measured as an inverse read-out of enzymatic product formation. Error bars represent standard deviation of initial slopes, individually calculated from 2 biological replicates, each with technical duplicates (n=4). Kcat values displayed, calculated from a non-linear regression curve fit (Vmax/FSP1 concentration [μM]).

**Figure S5.**
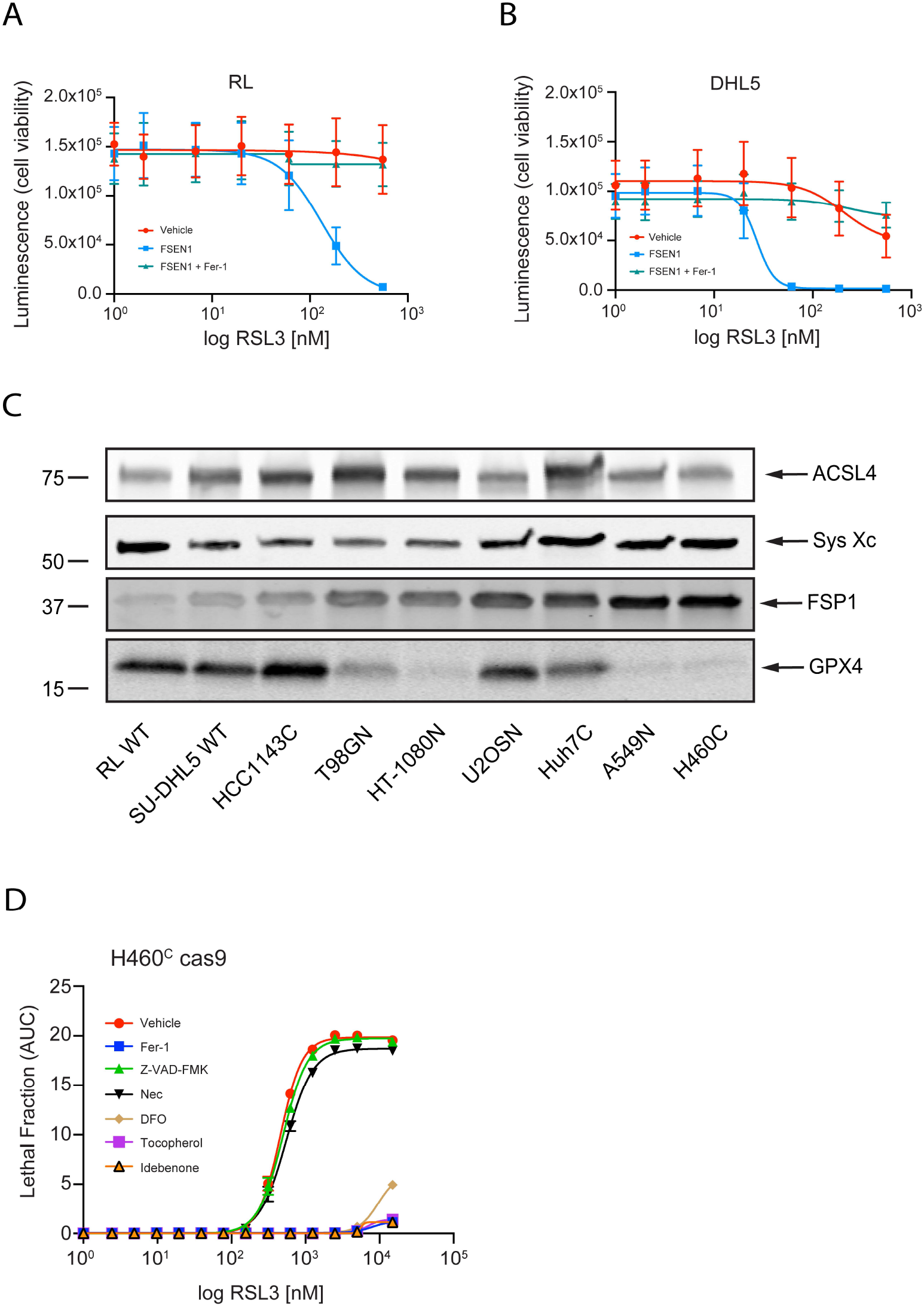
FSP1 inhibition triggers ferroptosis in cancer cell model. **A)** Dose response of RSL3 induced cell death in lymphoma cells 24 hr after co-treatment with vehicle or 1 µM FSEN1 or 1 µM FSEN1 together with 2 µM Fer-1 using the CellTiter-Glo Luminescent Cell Viability Assay. Data are mean ± SEM from three biological replicates. **B)** Western blot analysis of cell lysates from different cancer cell lines comparing SLC7A11, ACSL4, GPX4, and FSP1. **C)** Western blot analysis of cell lysates from different cancer cell lines comparing SLC7A11, ACSL4, GPX4, and FSP1. **D)** Dose response of RSL3-induced cell death in H460C Cas9 cells co-treated with 1 µM FSEN1 together with the indicated inhibitors of ferroptosis (Fer-1 [2 µM], DFO [100 µM], idebenone [10 µM], tococopherol [10 µM]), apoptosis (Z-VAD [10 µM]), and necroptosis (Nec1s [1 µM]). Data are mean ± SEM from three biological replicates. The Lethal Fraction (AUC) of 5 µM RSL3-induced cell death calculated from these experiments are also listed in **Figure 4**.

**Figure S6.**
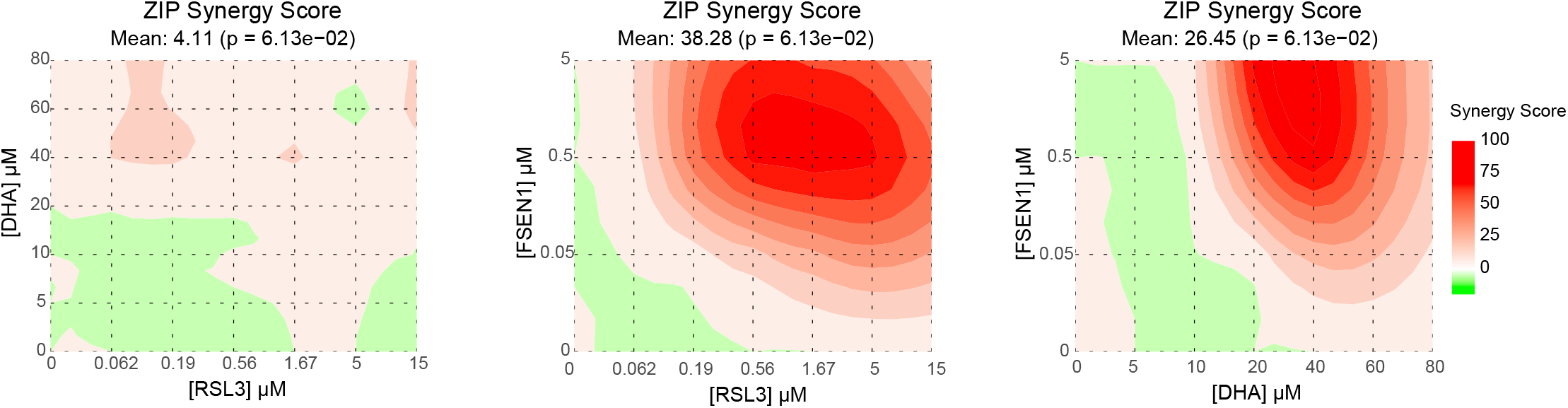
Drug contour maps. 2-D-dose response contour plots illustrating the distribution of synergy between FSEN1, DHA, and RSL3 in H460^C^ Cas9 cells, based upon **Figure 7C**.

